# Restriction-modification systems are required for *Neisseria gonorrhoeae* pilin antigenic variation

**DOI:** 10.64898/2026.01.10.698803

**Authors:** Selma Metaane, H Steven Seifert

## Abstract

*Neisseria gonorrhoeae* (Gc) pilin antigenic variation is a diversity-generating system that uses gene conversion to produce a variety of PilE protein variants, the major subunit of the Type IV pilus (T4p). Pilin antigenic variation allows the bacteria to escape immune surveillance and can alter T4p expression. While pilin antigenic variation requires many conserved homologous recombination and DNA repair factors, the pattern of sequence changes leading to pilin antigenic variants resembles that of an annealing reaction, rather than the expected long recombination tracts usually found in homologous recombination. We demonstrate that two paralogous restriction modification modules cleave specific, unmodified sequences within the expressed and silent pilin loci and that cleavage is essential for pilin antigenic variation. Moreover, these restriction activities affect the bacterium’s fitness. These findings partially explain the patchwork recombination patterns of pilin antigenic variants and suggest a unique mechanism for generating diversity.

## INTRODUCTION

*Neisseria gonorrhoeae* (Gc) is the causative agent of gonorrhea, a sexually transmitted infection that remains a global health concern due to its development of antimicrobial resistance ^1,2^. Gc is a human-restricted bacterium whose adhesion and invasion rely on pili and outer membrane proteins (reviewed in ^3^). Among them, the type IV pilus (T4p) is a critical virulence factor that facilitates bacterial colonization ^4–6^. Gc type IV pilin antigenic variation (Av) results in the existence of multiple versions of its major subunit, pilin (or PilE) ^7^ within a lineage. During pilin Av one of ∼19 silent copies (*pilS*) replaces the analogous sequences in the expressed *pilE* gene, while the *pilS* gene remains unchanged ^8,9^. This gene conversion process alters the PilE amino acids sequence, which generates a variable type IV pilus (T4p). Pilin Av can result in a fully expressed variant pilus, reduced T4p expression, or a completely nonpiliated cell ^10–12^. This extensive variability may promote persistence within the host, or more importantly, allow reinfection of a person who was previously colonized by the same bacterium ^13,14^.

The *pilE* gene itself consists of a conserved region followed by semi-variable and hyper-variable regions that share sequence identity with the *pilS* copies. There are two conserved elements called *cys1* and *cys2* surrounding the hypervariable loop that form a disulfide bond in pilin. The *pilE* ORF is flanked downstream by the *SmaCla* repeat, which is also found at the 3’ end of all *pilS* loci. The regions *cys1*, *cys2,* and *SmaCla* are involved in pilin Av ^15,16^. Pilin Av also involves transcription of the *gar* (G4-associated RNA) noncoding sRNA upstream of *pilE* to form a DNA:RNA hybrid (R-loop) ^17–19^. This R-loop captures the C-rich strand, enabling the formation of a guanine quadruplex (G4) structure ^17^. Mutation of either the *gar* promoter or the G4-forming sequence through single-base mutations abrogates pilin Av.

Pilin Av recombination tracts are defined by the first and last nucleotide changes that differ from the starting recipient *pilE* sequence, and these recombination tracks are flanked by regions of microhomology shared between *pilE* and the *pilS* donor ^12,20^. While pilin Av requires the *recA* gene ^21^, and the RecF-like pathway genes *recO*, *recR*, *recJ*, and *recQ* ^22–24^, the 10 to 500 bp regions of variant sequence transferred during pilin Av suggest that pilin AV requires other reactions besides homologous recombination. We have previously proposed that pilin Av involves two steps: an initial recombination between the *pilE* gene and a *pilS* copy at a region of microhomology, followed by homologous recombination of the hybrid *pilE-pilS* intermediate into an intact *pilE* ^20^.

Restriction-modification (RM) systems are widespread in prokaryotic genomes, as 83% of available genomes contain at least one of these modules ^25^. There are four major types of RMs ^26^, and type II RMs are the most abundant, accounting for 39.2% of bacterial genomes ^25^. Gc isolate FA1090 possesses 15 RM systems ^27–31^. The 11 FA1090 type II RM systems each include a sequence specific restriction endonuclease and a DNA methyltransferase. The endonuclease cleaves double-stranded DNA at its target motif unless the methyltransferase protects the site ^32–34^. Several host-restricted bacteria, such as *Helicobacter*, *Haemophilus*, *Streptococcus,* and *Neisseria,* all encode multiple RM modules, despite having relatively small genomes. Being host-restricted, we predict these organisms have fewer bacteriophage predators as compared to environmental bacteria; thus, it is unknown why these organisms have multiple RM modules ^35^. In this work, we demonstrate that specific 5’-CCGG sequences are frequently located at the microhomology bordering the recombinant pilin Av sequences. We show that the *pilE* and *pilS* 5’-CCGG sites, as well as the paralogous RM modules, are required for efficient pilin AV. Surprisingly, most genomic 5’-CCGG sequences are undermethylated and a subset are cut by the paralogous restriction endonucleases. Restriction of 5’CCGG sites in a subset of chromosomes results in a Gc fitness defect. These results indicate that Gc has co-opted these RM modules to facilitate pilin Av, albeit at the cost of reduced fitness.

## RESULTS

### A 5’-CCGG site is prevalent at the borders of pilE recombination tracts

Gc isolate FA1090 encodes 15 RM systems, which are conserved in most Gc isolates. Among all the putative RM systems present in FA1090, the recognition sites that are the most represented in the genome are 5’-GCSGC (16173 sites) and 5’-CCGG (11103 sites) (Table S1). Some sites overlap, such as 5’-G<u>CCGG</u>C, which also contains 5’-CCGG.

The 5’-CCGG motif shows a differential distribution across the Gc genome, with a frequency higher than one site per 100 bp in 167 out of the 2,211 annotated ORFs (Supplementary Table 3). All 19 *pilS* copies and the *pilE* variable region were among these high-density loci. In contrast, 5’-CCGG sites were completely absent from the *garP/G4* sequences and the *pilE* conserved N-terminal coding sequences (Fig. 1A, Table S2).

**Fig. 1.**
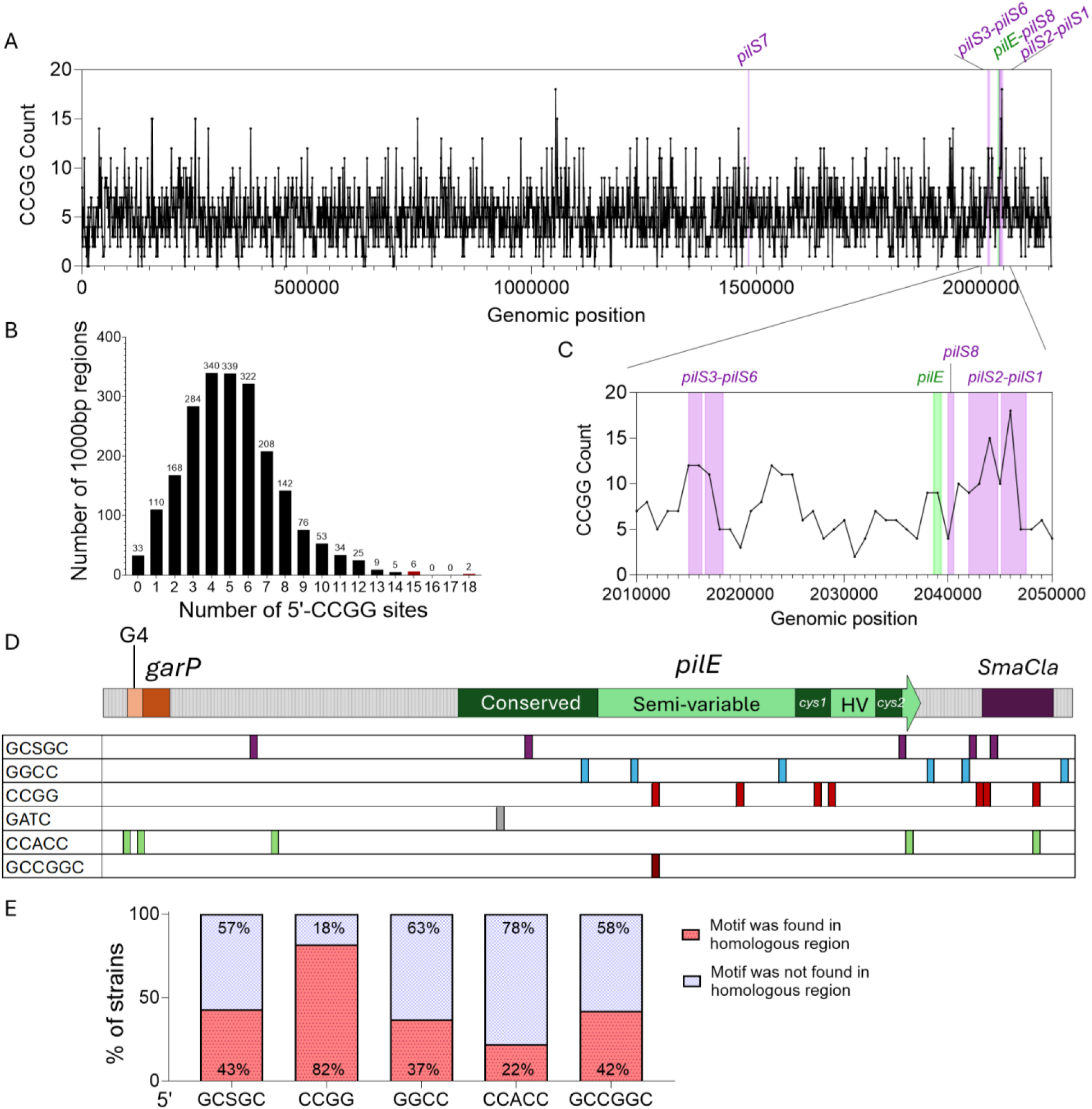
A restriction and modification target sequence is overrepresented in the *pilE gene.* **A.** Number of 5’-CCGG sites across Gc FA1090 genome per 1000 bp bin. **B**. Zoom-in on the *pilE* region from panel A. **C**. Distribution frequency of 5’-CCGG sites per 1000 bp bin across the genome. **D**. A schematic of the *pilE* locus. The cartoon shows the *garP* sRNA promoter, the guanine quadruplex-forming sequence (G4), the *pilE* open reading frame that includes the conserved N-terminus coding sequences and the semi-variable and hypervariable regions (that are also in the *pilS* copies), containing the conserved *cys1* and *cys2* repeats, followed by the *SmaCla* repeat. The *SmaCla* repeat is found at the end of each *pilS* locus. The 5’-CCGG is the most frequent in the *pilE* locus and is found only within the *pilE* variable region. **E.** Quantification of the RM target sites at pilin variant recombination tracts that were identified in the analysis of FA1090 pilin variants^12^. The 5’-CCGG motif is present at the site of recombination in 82% of the variant strains.

The pattern of variant *pilE* sequence changes has been extensively studied ^21,36–38^. We reanalyzed 100 previously described variant *pilE* sequences, identified their *pilS* donors, and mapped each recombination tract ^12^. The sequences bordering the recombination tracts upstream and downstream of the *pilS* insert indicate where the recombination event occurred. 82% of the analyzed *pilE* variants contained a 5’-CCGG site at one or both recombination track border (Fig. 1B). In contrast, the six *pilE* 5’-GGCC sites were only localized at the recombination tracts border of 37% of the variant strains (Fig 1A, B). Based on this reanalysis of published sequencing data, we postulate that the 5’-CCGG sequences could have a role in pilin Av.

### The pilE 5’-CCGG sites are important for pilin Av

There are several methods for measuring pilin Av frequencies ^22,39^. One method is the pilus-dependent colony morphology change (PDCMC) assay, which records the number of nonpiliated or underpiliated blebs that emerge from a piliated colony over time ^22^. When pilus phase variation occurs by pilin Av, they are captured by the PDCMC score ^22^. One limitation of this assay is that it is sensitive to growth rates since slower growth decreases antigenic variation ^39,40^. We introduced silent mutations that do not change the *pilE* coding sequence in all four 5’-CCGG sites (strain SM598). The PDCMC assay of strain SM598 with the mutated 5’-CCGG sites revealed a significant decrease in pilin Av (Fig. 2 B, E). After 30h of growth, the colonies of the *pilE* CCGG-disrupted mutant strain SM598 did not show any significant growth defect (Fig. 2A). Since the decrease in SM598 pilin Av frequencies is not due to decreased growth. These results support the hypothesis that the sites found at the borders of recombination tracts are important for pilin Av (Fig. 2A).

**Fig. 2.**
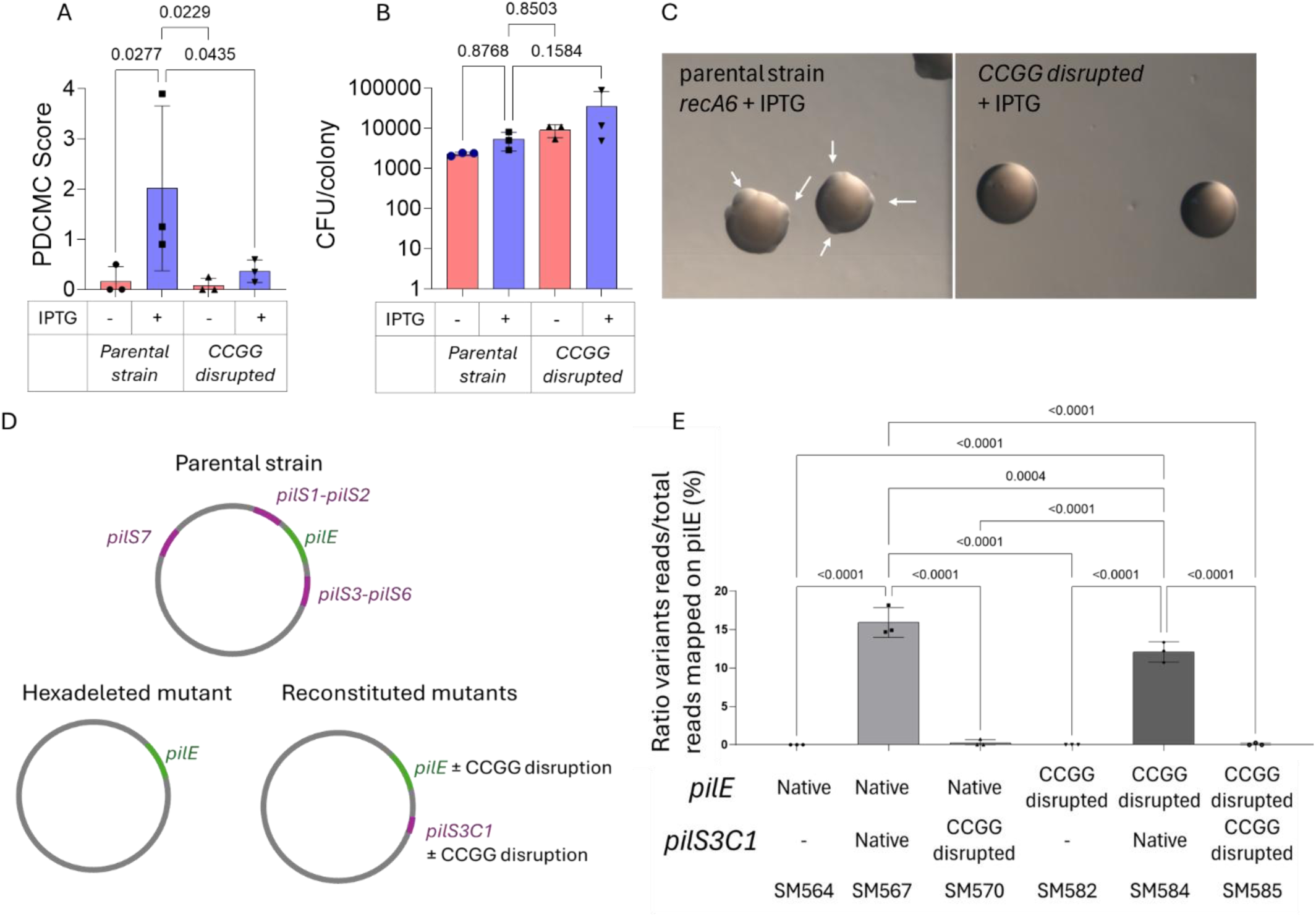
Pilin antigenic variation levels of the parental strain *recA6*. **A.** Pilin antigenic variation levels of *pilE* CCGG disrupted mutant (SM598) measured by the surrogate PDCMC assay in the absence or presence of 1 mM IPTG. **B.** Colony forming units (CFU) per colony at 30h of growth in the parental strain and the *pilE* CCGG-disrupted mutant (SM598) in the presence and absence of 1 mM IPTG**. C.** Pictures of colonies of the parental *recA6* strain and the *pilE* CCGG-disrupted mutant (SM598) after 32h of growth on GCB + IPTG. **D.** Schematic representation of the *pilE* (±CCGG disruption) and *pilS* (±CCGG disruption) constructs in a *pilS* hexadeleted mutant (SM564). **E.** PCR-based assay using Illumina sequencing to measure the variation level in a *pilE* amplicon. Approximately 500 colonies of the *pilS* hexadeleted strain were grown with 1mM IPTG. *pilE* PCR was performed, and Illumina sequencing of the native or mutated *pilE* and/or *pilS* loci. Fischer’s LSD uncorrected test was used; only p values <0.05 were plotted.

Despite the lower level of variation in the *pilE* 5’-CCGG mutant, some pilin variants were observed. Analysis of individually derived nonpiliated or underpiliated (P-) variants of SM598 revealed that 11/30 had a *pilE* gene deletion. Some P-variants (5/30) had the parental *pilE* sequence, indicating another process than pilin Av produced the nonpiliated phenotype. The remaining 14 variants were pilin antigenic variants. In these variant sequences, reconstituted 5’-CCGG sites were found in the *pilS* inserts, but not outside of the recombination tract. However, ten out of twelve *pilS* variant sequences contained 5’-CCGG sites at the variant border. Six had sites on both sides of the insert, while four had a site on one side. This analysis suggests that 5’-CCGG motifs in *pilS* may also contribute to pilin Av, even when they are no longer present in *pilE*.

### pilS copy 5’-CCGG Sites are critical for pilin Av

We tested whether the 5’-CCGG sites in the *pilS* donors are also involved in pilin Av. Given the presence of 19 *pilS* copies in the genome, mutating all 5’-CCGG sites within these loci proved challenging. Therefore, we generated a strain where all 19 *pilS* copies were deleted (the *pilS* hexadeleted mutant, SM564, Fig. 2D). We then reintroduced an unaltered *pilS3C1* copy into its original locus (Strain SM567) and a *pilS3C1* copy with five mutated 5’-CCGG sites (strain SM570). In these original and mutated *pilS3C1* strains, we also introduced the mutated *pilE* with or without 5’-CCGG sites (respectively, strains SM584 and SM585) (Fig. 2D).

Because the frequency of pilin Av is lower with a single *pilS* copy, we could not use the PDCMC assay to measure pilin Av. Instead, we used a modified PCR-based sequencing assay to measure pilin Av ^39,41^. The *pilS3C1-*mutated SM564 strain did not exhibit any detectable *pilE* variation (Fig. 2E), while the non-mutated *pilS3C1* strain SM567 showed typical pilin Av frequency (Fig. 2E). Mutating the *pilE* 5’-CCGG motifs with the nonmutated *pilS3C1* donor also reduced pilin Av (SM584). Finally, disruption of the 5’-CCGG motifs in both *pilE* and *pilS* reduced the level of antigenic variation to undetectable levels (SM585), identical to the hexadeleted control (SM564, Fig. 2E). These results show that 5’-CCGG sites in both the donor and recipient gene are critical for pilin Av.

### Two RM Modules targeting 5’-CCGG are required for Pilin Av

The pilin Av phenotypes of the *pilE* and *pilS* 5’-CCGG mutants suggested that these palindromic sites are recognized by factors that are critical for pilin Av. Gc encodes several RM systems ^27,42^. We identified three type II RM modules that are predicted to recognize 5’-CCGG and 5’-G<u>CCGG</u>C sequences. Two of these methylases were previously reported to methylate their predicted sites: m.NgoAXIV (RM1M, 5’-CCGG) and m.NgoAIV (RM3M, 5’-G<u>CCGG</u>C) ^27^. The m.NgoAXIII (RM2M) methylase is predicted to recognize 5’-CCGG ^42^, but this prediction has not been experimentally confirmed (Table S1). The *ngoAXIV* (RM1) operon encodes a restriction enzyme with two predicted ORFs (RM1R1 and RM1R2) and a complete methylase (RM1M), while the *ngoAXIII* (RM2) operon includes a full restriction enzyme (RM2R) and a methylase split into two predicted ORFs (RM2M1, RM2M2) (Fig. S2). Analysis of 47 complete genomes from PubMLST database shows that both operons are fully conserved across all analyzed Gc strains. Notably, 71% of *Neisseria meningitidis* genomes, a closely related species that also undergoes pilin Av, encoded a 5’-CCGG restriction enzyme (RM1R). In contrast, neither RM1R nor RM2R was detected in *Neisseria lactamica*, which does not undergo pilin Av (Supplementary Table 2).

To test the involvement of the RM modules in pilin Av, we constructed two mutant strains. We deleted the RM1 and RM2 operons that are predicted to recognize 5’-CCGG (strain SM500). We also deleted the RM3 operon that is predicted to recognize 5’-G<u>CCGG</u>C (strain SM604). The SM604 mutant showed no differences in growth or pilin Av frequency and was not analyzed further (Fig. 3B). In contrast, the RM1RM2 double mutant SM500 displayed a significantly lower level of PDCMC and increased growth (Fig. 3B, C). Moreover, deleting each restriction gene RM1R1 (SM323) and RM2R (SM327) also reduced pilin variation levels (Fig. 4A, B). Pilin Av was restored upon complementation of the RM1R1 or RM2R mutants with the mutated gene with a TetR-regulated promoter gene at an ectopic site (Fig. S3).

**Fig. 3.**
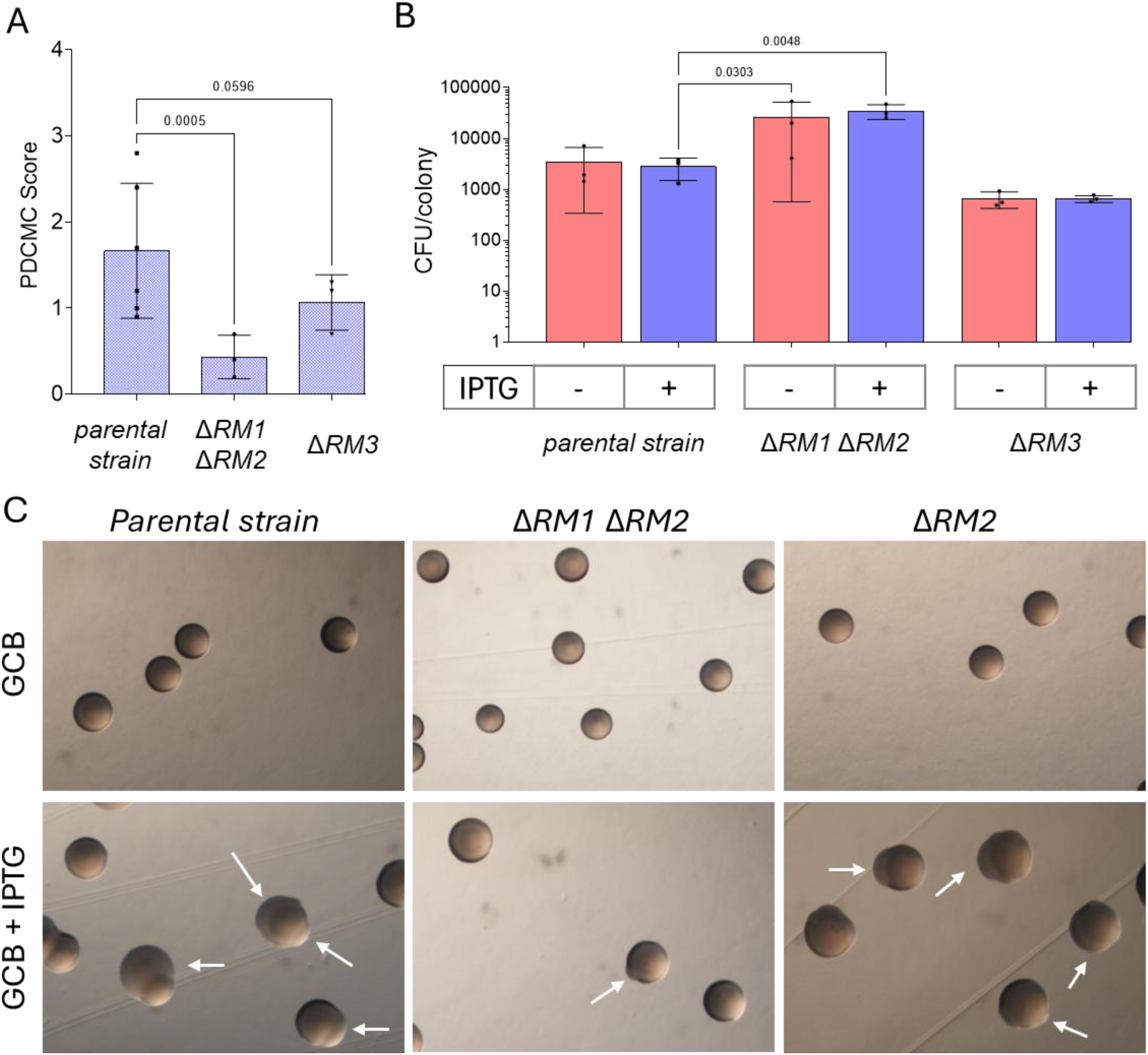
Mutation of RM genes alters pilin antigenic variation. **A.** Pilin Av levels of the parental strain and the SM500 and SM604 mutants measured by the PDCMC assay. **B.** Growth of the parental strain and the SM500 and SM604 mutants without IPTG on solid media. Fischer’s LSD test was used for multiple comparisons; only the p-values <0.05 were plotted. **C.** Pictures of the colonies grown with and without IPTG, showing the appearance of P-blebs (white arrows).

**Fig. 4.**
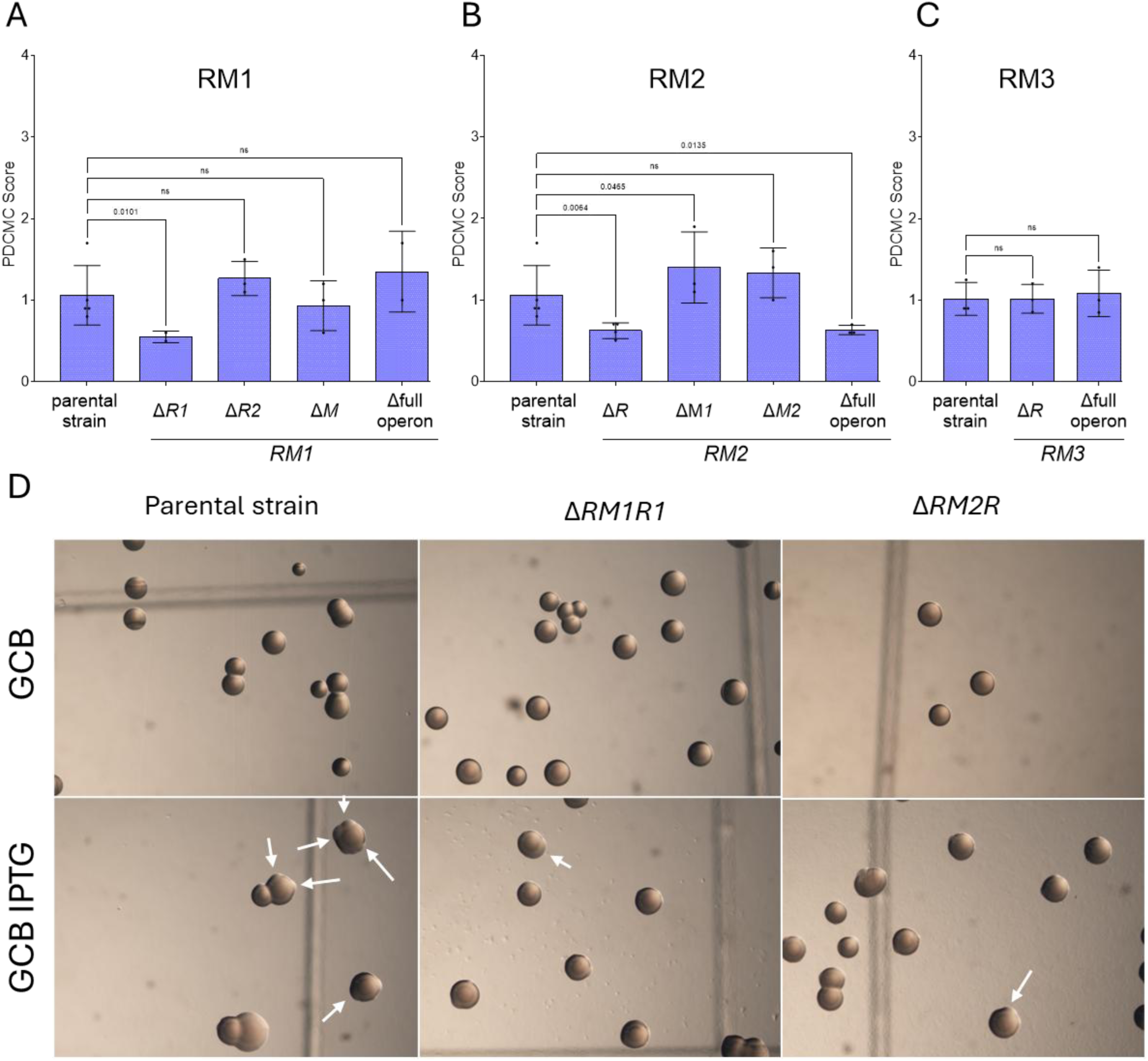
Deletion of RM system operons reduces pilin Av. PDCMC assay results (graphs A,B &C) and picture of the pilin Av defective colonies (D). A, B & C. Shown are the PDCMC scores for each single deleted mutant compared to the parental strain grown on GCB + IPTG for 30h. Multiple comparisons were performed using Fisher’s LSD test. D. Pictures of the colonies of the parental and the two mutants with a lower PDCMC than the parental, Δ*R1.RM1* and Δ*R.RM2* grown on GCB and GCB IPTG for 30h.

The observed reduction in PDCMC in the Δ*RM1R1* (SM323) and Δ*RM2R* (SM327) mutants was not due to a growth deficiency (Fig. S3). We also observed that the deletion of the entire RM2 operon results in similar levels of pilin Av to those of the RM2R mutant, whereas this was not the case for the RM1 operon (Fig. 3B, C). RNA sequencing data showed that the restriction gene RM2R has significantly higher transcript levels than its methylase^43^. Differences in transcription and stability between RM1 and RM2 effectors could explain why deleting the RM2 restriction enzyme has the same effect as deleting the whole operon. These results indicate that the restriction enzymes RM1R1 and RM2 are necessary for efficient pilin Av.

### The pilE and pilS 5’-CCGG motifs are cut in genomic DNA

To test the hypothesis that the restriction endonucleases RM1R1 and RM2R2 digest 5’-CCGG sequence, we designed an adapter-DNA with a GC overhang to ligate with the two nucleotide 5’-CG overhangs resulting from a 5’-CCGG cut. We ligated the adapter directly to purified genomic DNA, and PCR amplification was performed using a primer recognizing the adaptor paired with an upstream or downstream *pilE* primer (*pilRBS* or *pilEREV* primers ^44^). We detected multiple amplified products whose sizes corresponded to the location of *pilE* 5’-CCGG sites (Fig. S4).

We sequenced the adaptor-ligated DNA using ONT Nanopore long-read sequencing, which enabled us to distinguish the *pilE* and *pilS* loci. Among the four 5’-CCGG sites found in *pilE*, the adapter mapping revealed cleavage at the second and third 5’-CCGG sites. The first site, 5’-GCCGGC, and the fourth 5’-CCGG site (located 9 bp after the third site) showed no cuts (Fig. S6). In the *pilS* copies, cuts were detected at 44 of the 54 5’-CCGG sites across all 19 pilS copies, and no ligation was found at the 32 5’-GCCGGC sites. These cleavage patterns matched those observed in the positive control pretreated with HpaII, an endonuclease that also targets 5’-CCGG (Fig. 5, Supplementary Table 1). No cuts were detected in the RM1RM2-deleted strain (SM500), but cleavage was restored in the HpaII-treated control, confirming that deletion of these operons inactivates their cognate restriction enzymes. Notably, in the RM1RM2 double mutant pretreated with HpaII, we detected a low-level cut at the last 5’-CCGG site, suggesting that the residual loss of methylation upon RM1M and RM2M deletions implicates a minor protective role for at least one of the two 5’-CCGG-specific RM methylases (Fig. S5). As anticipated, the *pilE* CCGG-disrupted mutant (SM598) showed no cuts in *pilE*. However, all *pilS* had cleaved 5’-CCGG sites in SM598, as in the parental strain (Fig. 5, Supplementary Table 1).

**Fig. 5.**
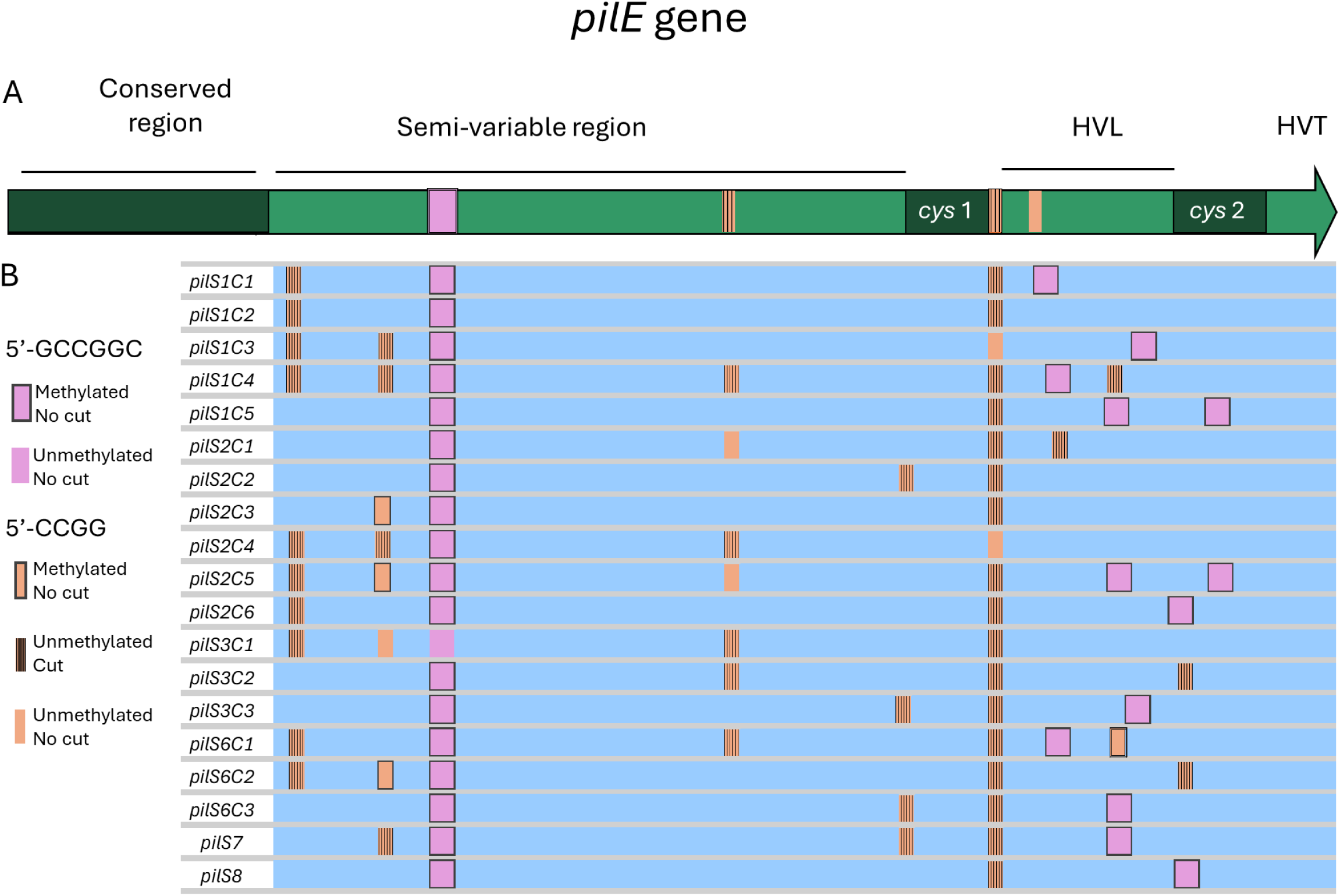
Mapping of the 5’-CCGG and 5’-GCCGGC cuts and methylation status within *pilE* and *pilS* copies. **A.** Schematic of the *pilE* locus with 5’-CCGG sites in orange and the 5’-GCCGGC site in pink. HVL corresponds to the hypervariable loop of *pilE* and HVT. **B** Schematic of the *19* aligned *pilS* with 5’-CCGG sites in orange and the 5’-GCCGGC site in pink. In A & C, the sites that exhibited cuts are hatched and the sites that were methylated are boxed. Sites are considered as methylated with a fraction N_modified_/N_valid_ > 3, and as cuts when the adapter was detected in more than 0.25% of the total reads at that genomic position.

### Methylation profiles of the pilE and pilS 5’-CCGG sites

Using nanopore sequencing, we quantified methylated residues (5mC) at each site with Modkit (Oxford Nanopore Technologies, 2023). Remarkably, across *pilE* and all *pilS* copies, most 5-GCCGGC were methylated (32/33 methylated sites), whereas most 5’-CCGG were not methylated (4/58 methylated sites) (Fig. 5 A,B). A more in-depth analysis of the methylated 5’CCGG showed these sites in particular overlap with other sites known to be a methylase target (5’-GGNNCC, NgoAXV; 5’-RGCGCY; NgoAI; 5’-GGCC, NgoAII). We found no cuts at methylated 5’-GCCGGC sites, confirming their protection. Similarly, methylated 5’-CCGG sites remained intact, and we didn’t detect methylation at the 5’-CCGG sites that were cleaved. Only five *pilE* 5’-CCGG sites were neither methylated nor cut. For this analysis, a cut was defined as detectable if it occurred in ≥0.25% of the reads at a specific site. In general, the 5’-CCGG sites are found to be undermethylated on the whole genome (Supplementary Table 4). This low level of genome-wide methylation at 5’-CCGG sites is consistent with PacBio data for Gc FA1090 deposited to the Rebase database ^42^. Additionally, the RM2 methylase (M.NgoAXIII) is predicted to target 5’-CCGG sites as an isoschizomer of HpaIIM, previous studies were unable to identify its target site based on methylation data and concluded that this methylase is inactive ^27^. Taken together, these results show that there is an undermethylation of some but not all sites within the genome.

### Restriction at 5’-CCGG sites impacts Gc growth

The RM1RM2 double mutant (SM500) displayed an enhanced growth phenotype after 30h of growth (Fig. 3A). The *pilE* CCGG-disrupted mutant (SM598) also displayed bigger colonies than the parental strain, although difference in CFUs was not significant (Fig. 2A). Since reduced growth can equalize after extended growth, we measured the CFU/colony of these strains during a growth curve on solid media. This solid media growth curve confirmed that SM598 showed enhanced growth compared to the parental strain (Fig. 6 C, D). In addition, strains lacking the 5’-CCGG-targeting restriction enzymes (RM1R1 and RM2R) also showed significantly higher CFU/colony counts, and growth was reduced by expressing RM1R or RM2R from an ectopic locus (Fig. 6D, F).

**Fig. 6.**
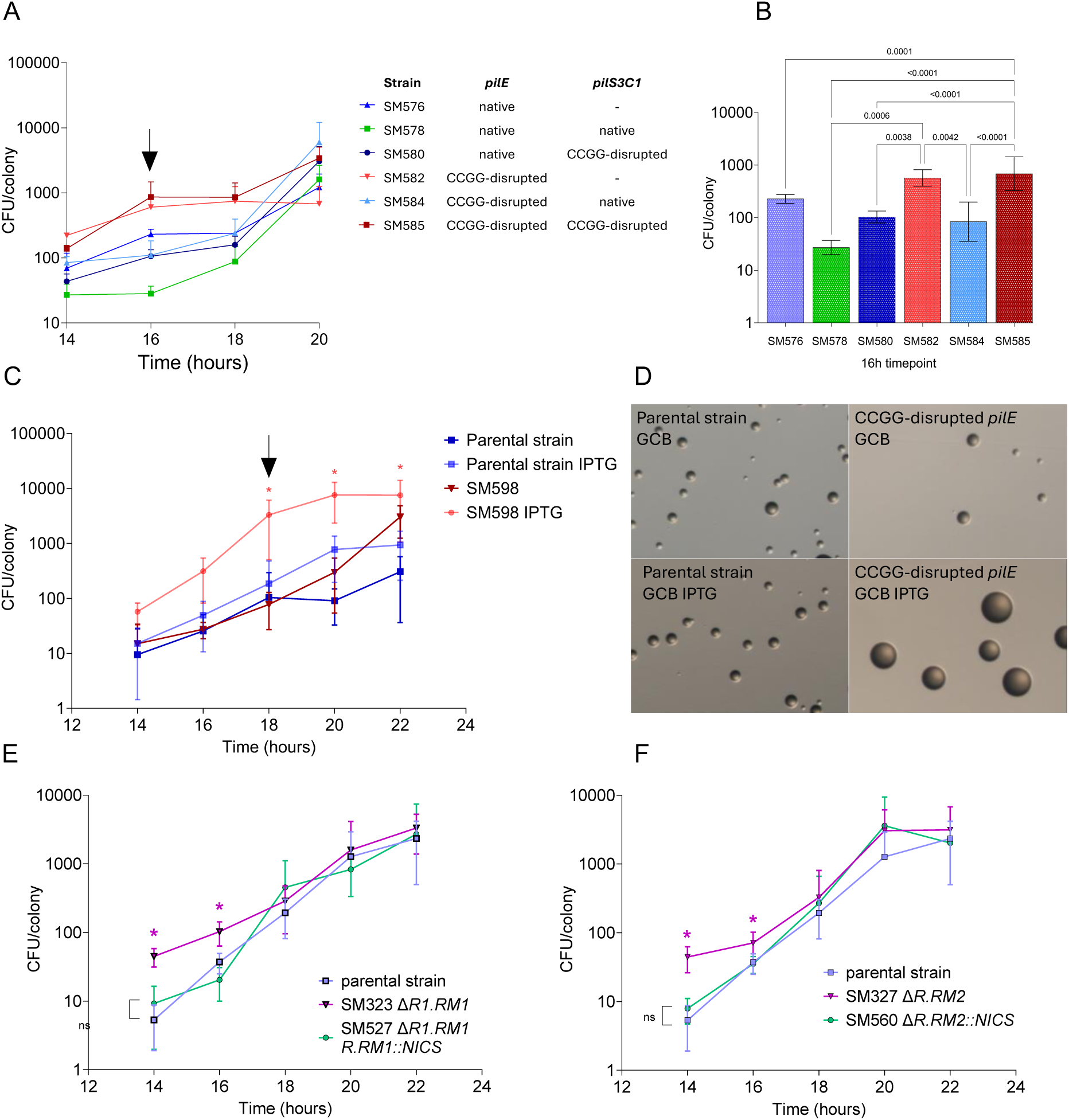
Growth on solid media. **A.** Strains from the *pilS* hexadeleted background with and without disruption of the *pilE* or *pilS1C3* sequences grown in the presence of 1mM IPTG. **B.** Statistical analysis using Fisher’s LSD multiple comparison of the 16h timepoint from panel A, only statistically significant comparisons are plotted. **C.** Growth of the parental strain and SM598 (*pilE* CCGG disrupted) in the absence or presence of 1mM IPTG. **D.** Pictures of the parental strain and the SM598 strain (*pilE* CCGG disrupted) grown for 18h on GCB in the absence or presence of 1mM IPTG. **E.** Complementation of the SM323 growth in the presence or absence of 1mM IPTG. F. Complementation of the SM327 growth in the presence or absence of 1mM IPTG.

We measured CFU/colony in hexadeleted strains carrying native pilin loci and the reintroduced *pilS* gene, with and without 5’-CCGG-mutation (Fig. 2A, B). Interestingly, the two strains that showed the highest CFU/colony were the *pilE* CCGG-disrupted mutant lacking *pilS* (SM582) or with a 5’-CCGG mutant *pilS* locus (SM585). The strain with native *pilE* and *pilS* sequences (SM578) demonstrated the lowest growth (Fig. 6A). Strains with either an intact *pilS* or *pilE* locus showed intermediate growth, confirming that the elevated CFU/colony count in the *pilE* 5’-CCGG-disrupted mutant is attributable to *pilE* and *pilS* disruption.

Collectively, these data indicate that RM1R or RM2R activity at 5’-CCGG sites in *pilS* or *pilE* results in reduced growth. This observation explains why unpiliated colonies tend to have higher CFU counts ^46^. To test this hypothesis, we compared the CFU per colony between two nonpiliated mutants: a Δ*pilE* mutant, which does not undergo pilin Av, and a *pilE* promoter mutant, a mutation that doesn’t affect pilin Av frequency ^47^. Interestingly, the *pilE* promoter mutation did not alter CFU per colony compared to the parental strain, whereas the Δ*pilE* mutant exhibited a significantly higher CFU per colony count (Fig. S7). We introduced the *pilE* promoter mutation in the CCGG-disrupted SM598 mutant, but it didn’t affect growth. This result shows that the increased growth we observed is most likely due to the CCGG mutation rather than pilin or pilus expression. These results suggest that cleavage of the *pilE* and *pilS* copies, and, by extension, pilin Av, negatively impacts fitness.

## DISCUSSION

Bacterial adaptation is shaped by a fundamental balance: traits such as virulence factors, antibiotic tolerance, or antigenic variation that may reduce individual fitness but allow populations to overcome selective pressures – whether from host immunity, microbial competition, or environmental challenges. Antigenic variation (Av) illustrates this paradox: by generating diverse surface antigen repertoires, some cells escape immune detection, while those with less variation are more likely to be eliminated ^48,49^.

We demonstrate that the paralogous restriction enzymes RM1R1 (R1.NgoAXIV) and RM2R (R.NgoAXIII), which each cleave the unmethylated 5’-CCGG sites across the genome to induce cuts within the *pilE* and *pilS* loci. We propose that these cuts ligate to create one or two junctions between the recombining genes, and resulting in regions of microhomology at one or both recombination junctions. These results help explain why small regions of DNA can be transferred during the pilin Av gene conversion reactions. However, these results do not explain the requirement for homologous recombination (e.g., RecA, RecO, and RecR) for pilin AV. Previous data suggested that there is a multi-step process in which a junction is formed between *pilE* and a *pilS* copy ^20^. We postulate that the hybrid locus forms on a donor chromosome and recombines with the *pilE* on the recipient chromosome ^20,50^. This idea is supported by the fact that Gc and the closely related *N. meningitidis,* both of which undergo pilin Av ^51,52^ are polyploid, while *N. lactamica* that does not undergo pilin Av is monoploid ^50^. This model is consistent with the observation that some *pilE* recombinants have only one 5’-CCGG site at the recombination tract border. Ultimately, invoking separate donor and recipient chromosomes explains why this represents a gene conversion process since the silent copies on the recipient chromosome are not involved. Notably, *vlsE* antigenic variation in *Borrelia* species, which are also polyploid ^53^ and possess several RM systems on their plasmids ^42^, occurs via *trans* recombination, a process in which genetic exchange takes place between physically distinct DNA molecules ^54^. It is interesting to speculate that *Borrelia* may also use RM system-mediated recombination between donor and recipient sequences as a step during antigenic variation. One unexpected observation from this study is the increased growth in RM mutants or strains with mutated *pilE* or *pilS* 5’-CCGG sites. These results suggest that restriction in the *pilE* and *pilS* 5’-CCGG sites interferes with Gc fitness. We knew that a Δ*pilE* mutant exhibits increased growth, and we assumed this was a fitness cost of producing the pilin protein and pilus fiber. However, since a *pilE* promoter -10 mutant (which abolishes *pilE* expression and thus piliation) displays parental-growth, we conclude that the reduced growth when the 5’-CCGG sites are cleaved is due to chromosomal loss during pilin AV.

There are many unknown parts of the pilin Av process. Our results do not explain how the digested *pilE* and *pilS* copies associate, whether multiple digestions events occur within the pilin loci on a single chromosome, and if other parts of the digested chromosomes would ligate together. Given that most pilin loci are clustered within a 30 kb region of the 2.1 Mb chromosome, hybrid formation may simply result from the proximity of these loci and their high density of 5’-CCGG sites. While we postulate that recombination occurs between a hybrid formed between a donor chromosome and a recipient, this would require protection of the second chromosome from digestion. Alternatively, hybrid locus formation could occur in one half of a diplococcus, with the recipient chromosome residing in the second coccal unit.

Taken together, these results imply that RM-mediated cleavage at 5’-CCGG sites in *pilE* is an early and critical step in pilin Av. This improvement in fitness in laboratory conditions was inversely correlated with the decrease in Pilin Av frequencies, which implies that pilin Av has a fitness cost even without immune pressure. This aligns with observations in other pathogens, such as malaria parasites and viruses, where antigenic diversity involves fitness costs *in vitro* ^55,56^. These findings advocate that the pilin Av mechanism is adapted to limit the proliferation of non-varying cells, thereby maintaining population-level diversity. These observations reinforce a key evolutionary principle: population resilience often depends on limiting individual fitness.

## METHODS

### Bacterial strains and growth

Bacterial strains used in this study (listed in Table S3) are derivatives of the *N. gonorrhoeae* FA1090 isolate. All strains were checked to have the *pilE* 1-81-S2 variant sequence developed during human volunteer colonization ^44^. The strains were grown in 37°C at 5% CO_2_ on Gonococcal (Gc) Medium Base (BD Difco) (36.25 g/L), agar (1.25g/L), Kellogg Supplement I (22.2 mM glucose, 0.68 mM glutamine, 0.45 mM cocarboxylase), and Kellogg Supplement II [1.23 µM Fe(NO_3_)_3_] ^11^.

### Pilin antigenic variation measurements

#### PDCMC pilin Av assay

The PDCMC assay was performed as previously published ^22^. The Gc strains were revived on GCB from a frozen stock overnight. The next day, a single colony was reisolated onto GCB and GCB 1 mM IPTG plates and incubated overnight. After 18h of growth, 20 colonies were selected, and their morphologies were monitored to score the appearance of nonpiliated blebs. After 30h of growth, the number of blebs in the selected colonies was counted. The variation score was counted e.g., 1 for 1 bleb, 2 for 2 blebs, 3 for 3 blebs, and 4 for 4 or more blebs. The PDCMC score corresponds to this variation score divided by the number of colonies (20).

#### Analysis of pilE variants

After 30h of growth on GCB 1mM IPTG, a bleb was reisolated from each colony on GCB. The next day, a single colony was used to perform a PCR with the primers PilRBS and S3PA ^44^. The amplicon was then sequenced using Genewiz Azenta Sanger services. The alignment of the sequences was performed using Jalview ^57^.

#### Pilin Av DNA sequencing assay

To enable measuring of pilin Av frequencies by Illumina sequencing, *recA6* strains were grown for 22 h on 1 mM IPTG GCB, which corresponds to about 19 or 20 generations ^39^. About 500 colonies were collected, and genomic DNA was extracted. The KOD Hot Start (Novagen, Toyobo) PCR was performed according to the supplied protocol, with 100 ng of template genomic DNA. False “variants” can occur by *in vitro* recombination during the PCR amplification ^39^. To limit these PCR-generated hybrids, we limited the number of cycles to 30. The PCR product was then purified using Qiagen PCR purification kit, and Illumina sequencing (SeqCenter). The sequencing data were aligned with *pilE* 1-81-S2 sequence as a reference genome using Hisat2 ^58^ and Samtools ^59^. The variant calling was then run using Pysam ^60^ in a custom script ^61^.

### Detection of restricted 5’-CCGG sites

A synthetic adaptor was produced with a 5’-GC overhang and a single phosphorylated 5’-end. To prepare the adapter-ligated DNA, we annealed the oligonucleotides SM243 TOP (CGGGTCGGCAGTAACGTATTGATGCATACC) and SM244 BOT (GGTCGGCAGTAACGTATTGATGCATACC<u>CG</u>) to create a CG overhang by mixing SM244 BOT and SM243 TOP, heating the mixture to 98°C, and gradually decreasing the temperature by 5°C every 10 minutes until reaching 4°C to allow efficient annealing. We purified the product using the column cleanup PCR purification kit (Qiagen). We ligated the purified adapters with purified SM18 *recA6* genomic DNA using T4 DNA ligase at 16°C overnight (NEB). For the positive control of this experiment, we used genomic DNA pre-treated with the restriction endonuclease Hpa II (NEB). PCR was performed using forward and reversed primers corresponding to the adapter sequences and the primers pilRBS ^44^, *pilE*REV ^44^, SM241 (GGCGTTACCGCGCCCGTC) and SM242 (ATCCATGAACCCGACCGCACAACG). The reaction was performed using GoTaq (Promega) with the adapter-ligated gDNA as the template. The amplified products were analyzed on agarose gels.

We performed nanopore sequencing of the adapter-ligated gDNA (SeqCenter). We analyzed the sequencing data using a custom script that mapped the adapters across the genome ^61^. The script processes nanopore sequencing data to identify adapter positions by mapping the FASTQ reads to a reference genome using minimap2, detecting adapters with edlib, and splitting reads at adapter sites. Adapter positions were filtered and the location of adaptors was visualized using matplotlib. The workflow utilizes pysam for BAM/FASTA file handling and Bio.Seq for sequence manipulation, providing a comprehensive analysis of sequencing data quality and biological significance.

### Methylation analysis

Methylated bases were identified using Modkit dist_modkit_v0.4.4_7cf558c ^45^, a tool designed for analyzing nanopore sequencing data. Raw sequencing reads were aligned to the reference genome using EPI2ME with default parameters. The resulting BAM files were processed with Modkit module for detection of the 5mC methylations. Methylation frequencies were visualized, and regions of interest were extracted for downstream analysis, comparing treated and control samples.

## QUANTIFICATION AND STATISTICAL ANALYSIS

The statistical analysis was conducted using GraphPad Prism. The growth and PDCMC statistical analysis were executed using an ANOVA test with the multiple comparison Fisher’s LSD parameters. Each bar graph represents the individual values that are represented by symbols. The bar graphs correspond to the mean, with the SD indicated as an error bar.

## ACKNOWLEDGEMENTS

We would like to thank Dr. Shaohui Yin for providing the hexadeleted mutant. We are also grateful to Dr. Linda Hu for her insightful and constructive feedback during the revision of this manuscript, and to all the members of the Seifert lab for their valuable discussions throughout the study.

Research reported in this publication was supported by the National Institute of Allergy and Infectious Disease (NIAID) of the National Institutes of Health under grant numbers R37 AI033493, R01AI146073, and R21AI148981.

## DETAILS

The content is solely the responsibility of the authors and does not necessarily represent the official views of the National Institutes of Health. This manuscript is the result of funding in whole or in part by the National Institutes of Health (NIH). It is subject to the NIH Public Access Policy. Through acceptance of this federal funding, NIH has been given a right to make this manuscript publicly available in PubMed Central upon the Official Date of Publication, as defined by NIH.

## EXTENDED DATA

**Table S1.** List of the predicted Type II RM with their predicted recognition motif and the number of occurrences in the Gc strain FA1090 N-1-60.

| Predicted Type II RM system | Motif (5' to 3') | Number of occurrences in FA1090 N-1-60 | Number of non-overlapping occurrences in FA1090 N-1-60 |
| --- | --- | --- | --- |
| NgoFVII | GCSGC | 16173 | 13952 |
| NgoAXIV and NgoAXIII | CCGG | 11103 | 7884 |
| NgoAII | GGCC | 4590 | 2018 |
| NgoAXIP | GATC | 2259 | 2112 |
| NgoAXV | GGNNCC | 2033 | 818 |
| NgoAX | CCACC | 1575 | 1380 |
| NgoAIV | GCCGGC | 1572 | 0 |
| NgoAI | RGCGCY | 622 | 224 |
| NgoAVIII | GACNNNNNTGA | 407 | 338 |
| NgoAIII | CCGCGG | 218 | 45 |
| NgoAXVII | GAGNNNNNTAC | 118 | 89 |

**Table S2.** Number of occurrences of each of the known motifs targeted by type II restriction-modification systems in Gc strain FA1090.

| Motif (5' to 3') | # in *pilE* | # in 19 *pilS* copies |
| --- | --- | --- |
| GCSGC | 2 | 41 |
| CCGG | 4 | 87 |
| GGCC | 3 | 48 |
| GATC | 1 | 1 |
| GGNNCC | 0 | 14 |
| CCACC | 1 | 12 |
| GCCGGC | 1 | 32 |
| RGCGCY | 0 | 8 |
| GACNNNNNTGA | 0 | 1 |
| CCGCGG | 0 | 0 |
| GAGNNNNNTAC | 0 | 0 |

**Fig. S1.**
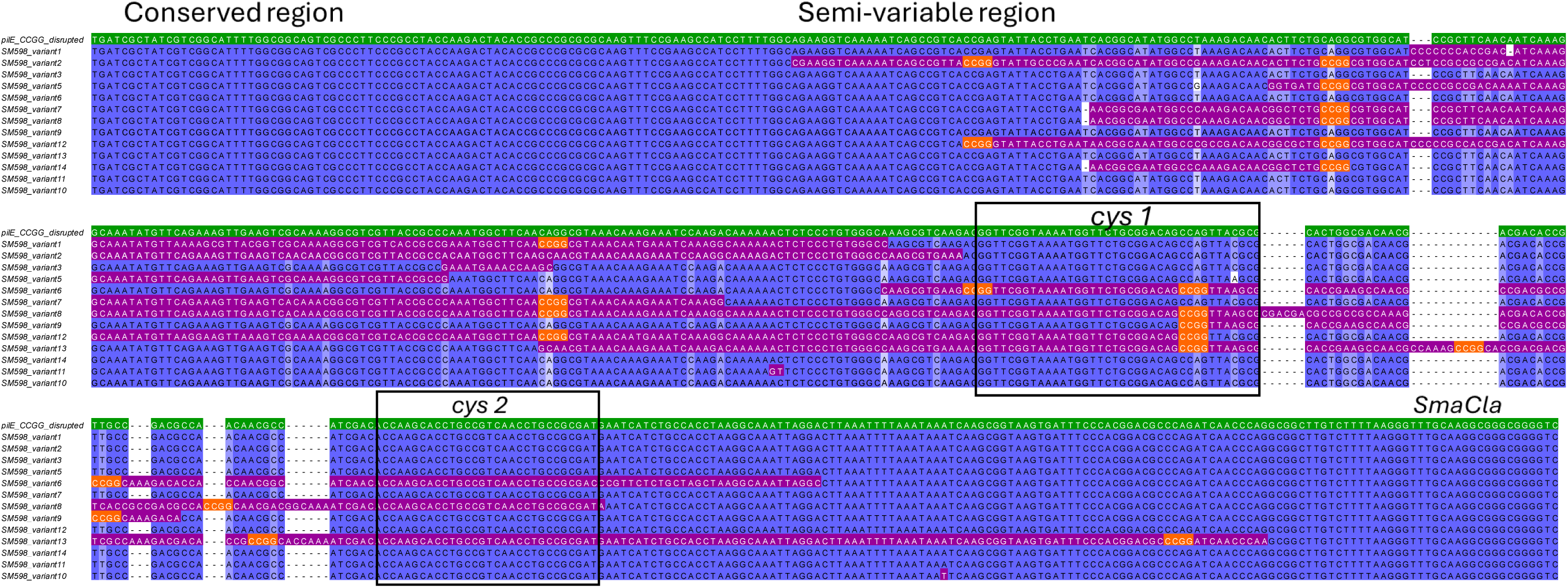
Sequence alignment of *pilE* amplicon from the SM598 mutant variants. The variants were reisolated from blebs on GCB IPTG after 30h of growth. The reference sequence is indicated in green, the *pilS* inserts are indicated in purple and the 5’-CCGG sites are indicated in orange.

**Fig. S2.**
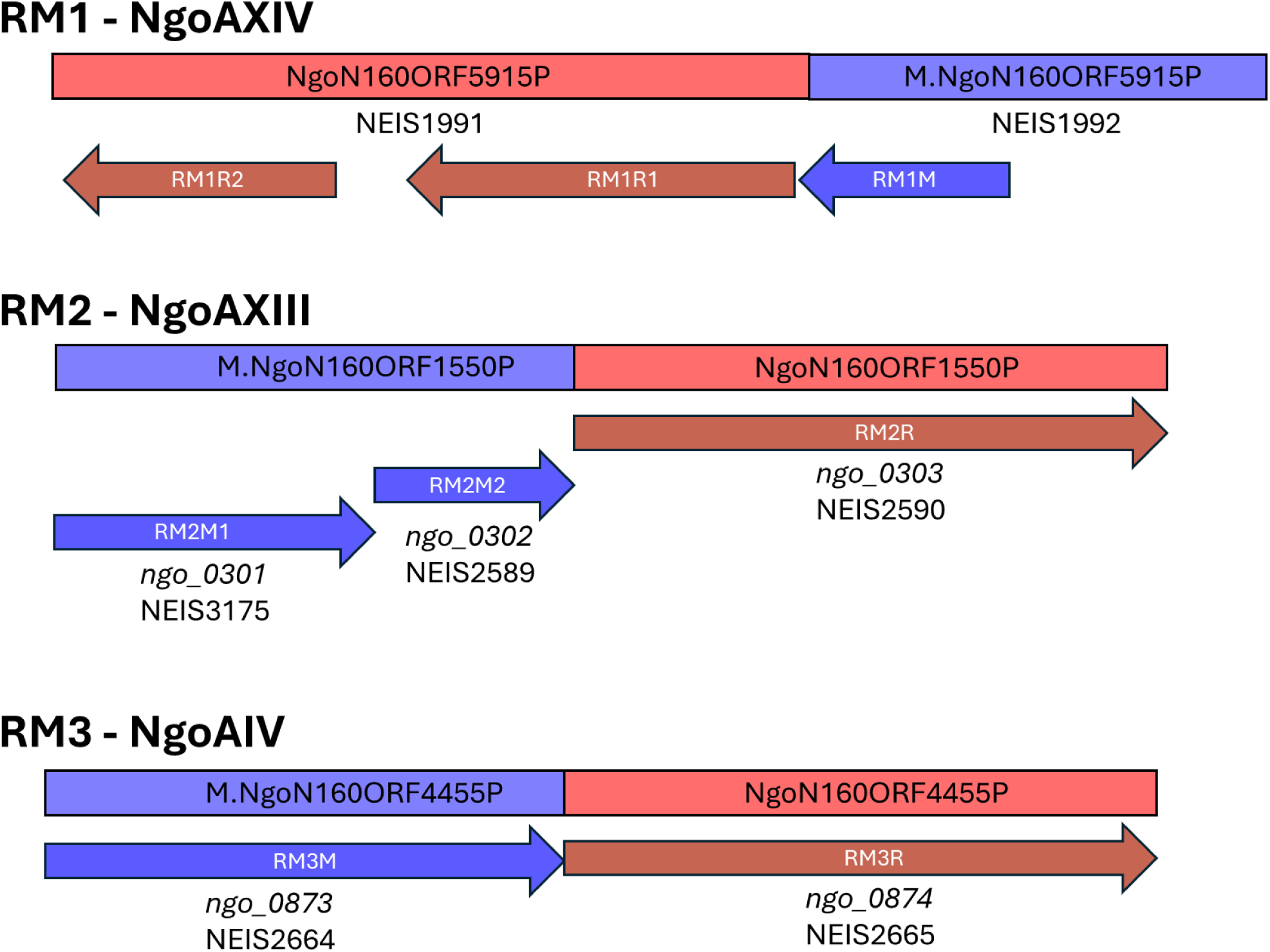
Genomic organization of the RM1 RM2 and RM3 operons in Gc FA1090. Rectangles follow Rebase nomenclature. ORFs are labeled with Ngo_ identifiers, while NEIS numbers corresponds to PubMLST nomenclature. Red indicates restriction genes; blue, methylase genes.

**Fig. S3.**
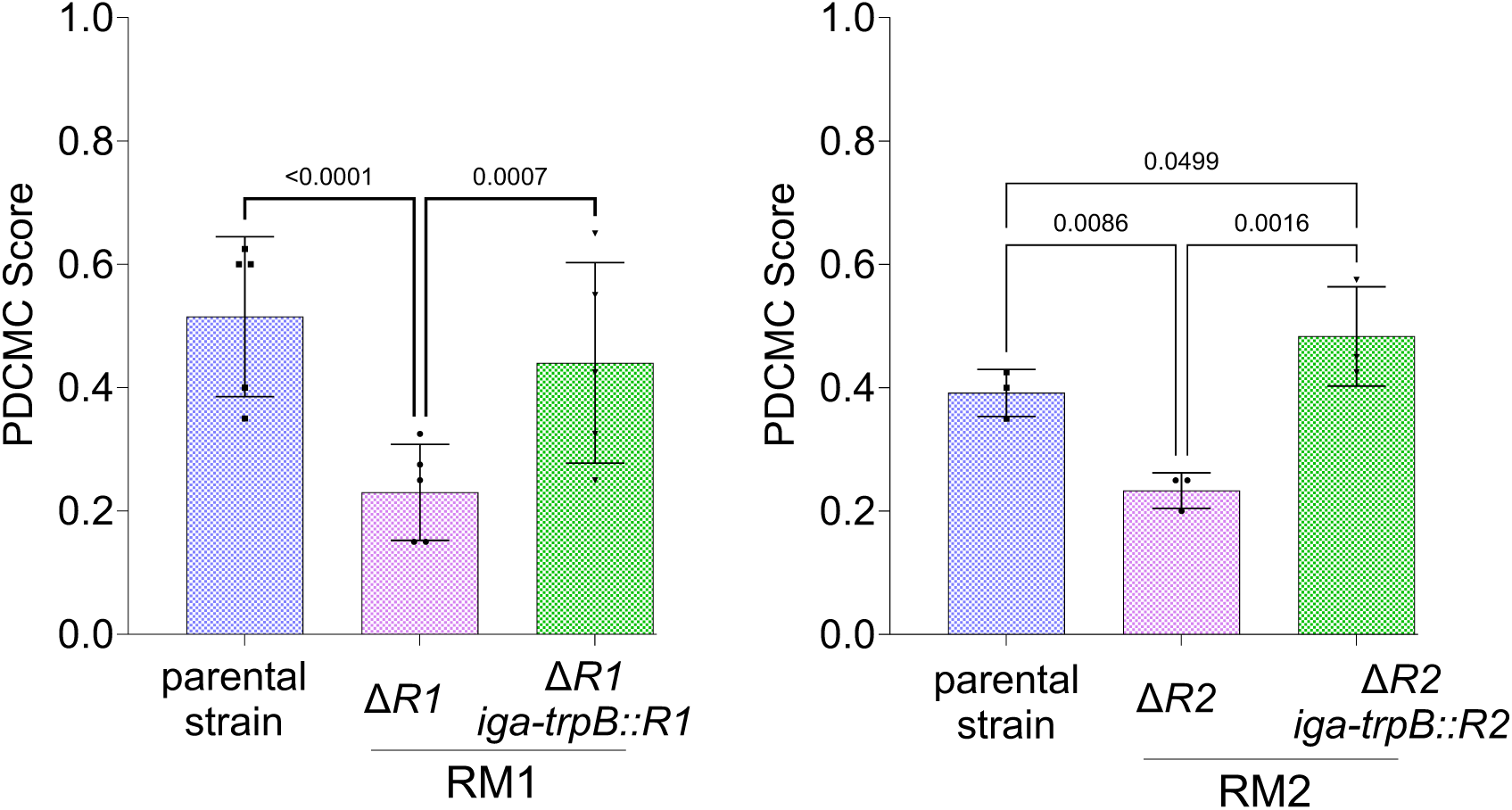
PDCMC score measured in the strains Δ*R1.RM1* (SM323) and Δ*R.RM2* (SM327) and their complements. The gene of interest RMR1 on the left, RM2R on the right was reinserted in the *iga-trpB* site using pMR69. Fischer’s LSD test was used, only the p-values <0.05 were plotted.

**Fig. S4.**
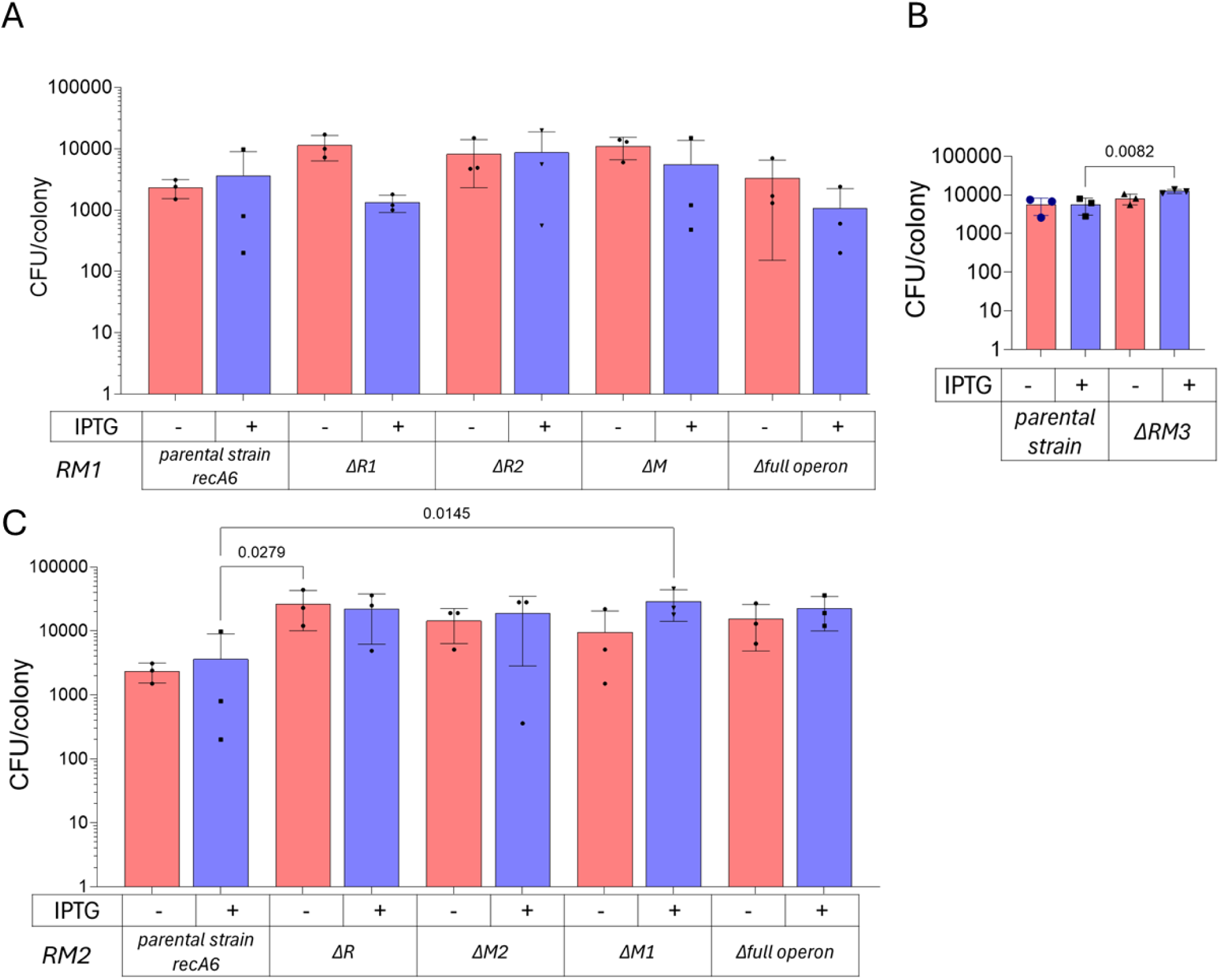
Measure of the growth of mutants carrying single gene deletions of the three operons. CFU/colony of the strains of the individual mutants for the operons **A.** RM1, **B.** RM3, and **C**. RM2, after 30h of growth on GCB with and without IPTG. Multiple comparisons were performed using Fischer’s LSD test, only the p-values <0.05 were plotted.

**Fig. S5.**
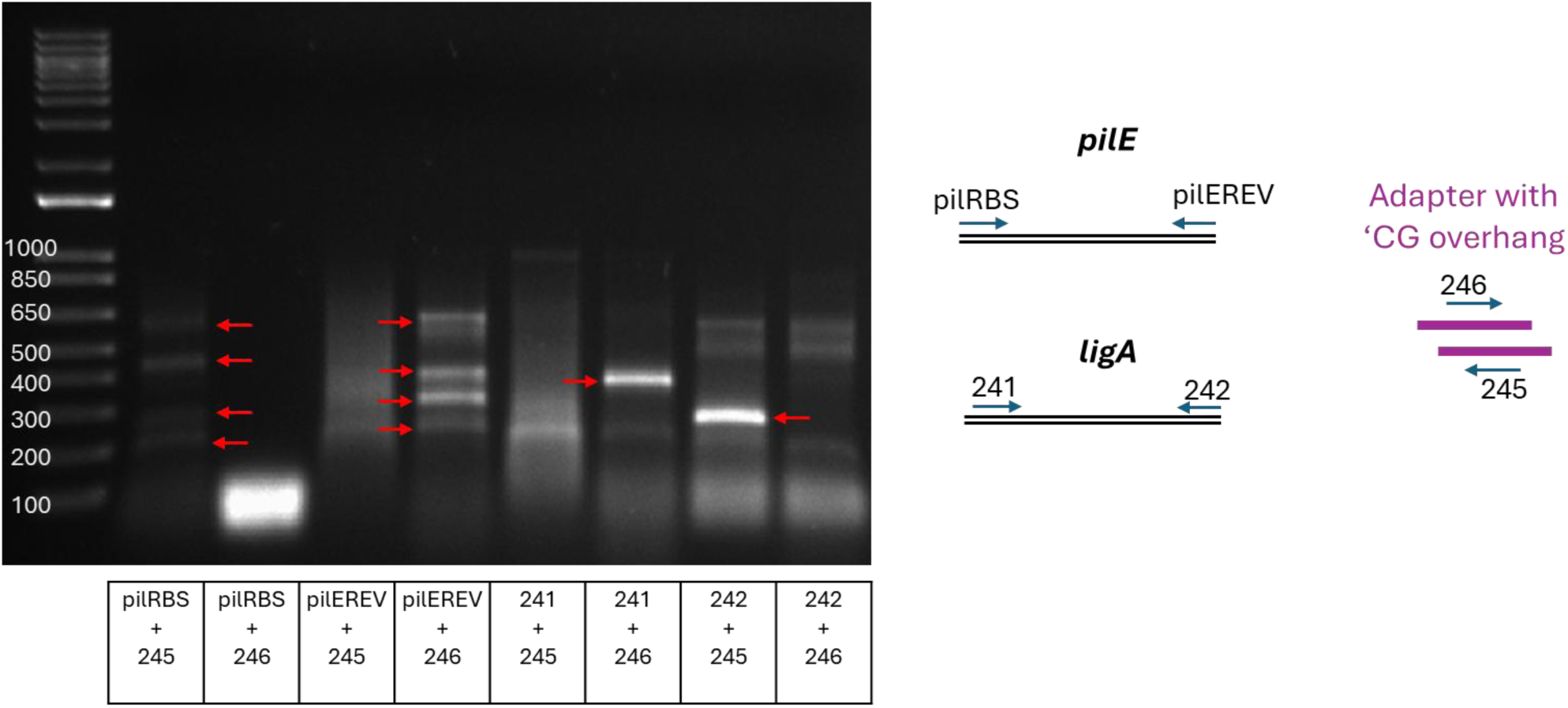
Ligation of the adapter within the genomic DNA. Migration of the PCR-amplified fragments using ligated genomic DNA as a template and several pairs of primers where one of them is internal to the ligated adapter (246 or 245). The bands expected if the adapter has ligated into a 5’-CCGG cut are boxed in reindicated with red arrows. The *ligA* gene is used as a control fragment that contains only one 5’-CCGG site.

**Fig. S6.**
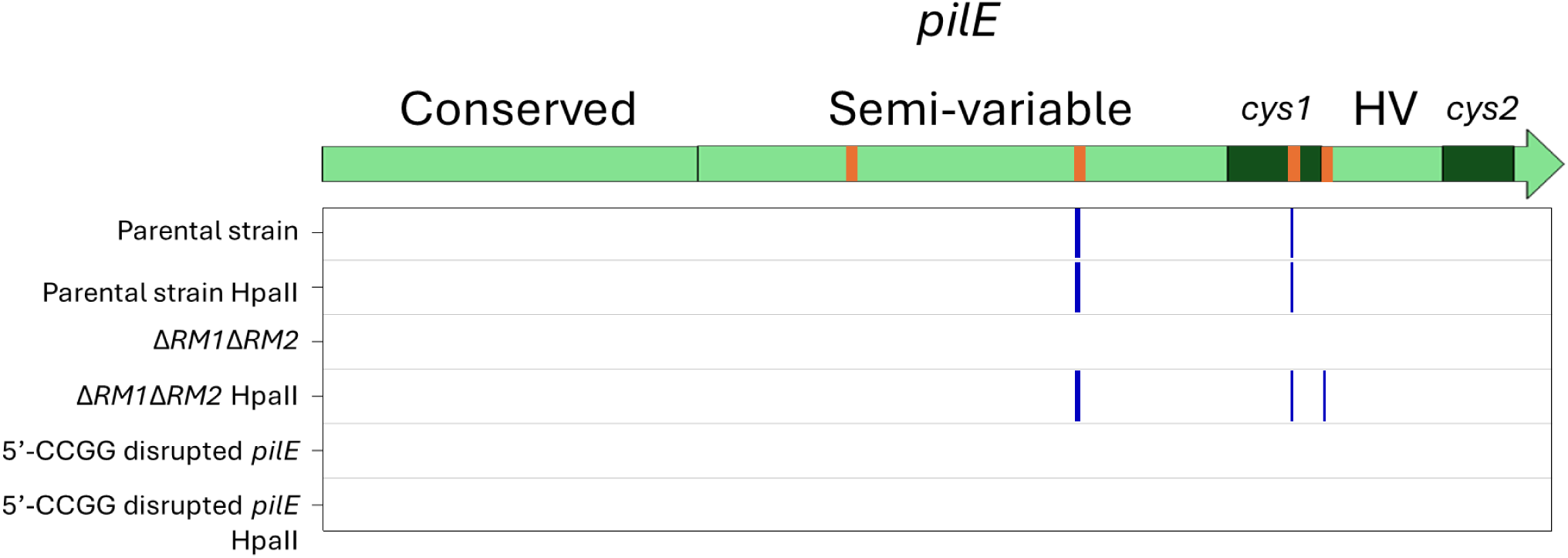
Detecting CCGG cuts within the *pilE* gene. Detection of the adapter within *pilE*. The schematic representation of the *pilE* shows its different regions with the 5’-CCGG sites are represented in orange. The panel below the *pilE* gene shows the locations of the detected CCGG cuts in, from top to bottom: the parental strain *recA6*, it’s HpaII-pretreated control, followed with the RM1RM2 double mutant (SM500), its HpaII control, the CCGG-disrupted strain (SM598) and its HpaII control.

**Fig. S7.**
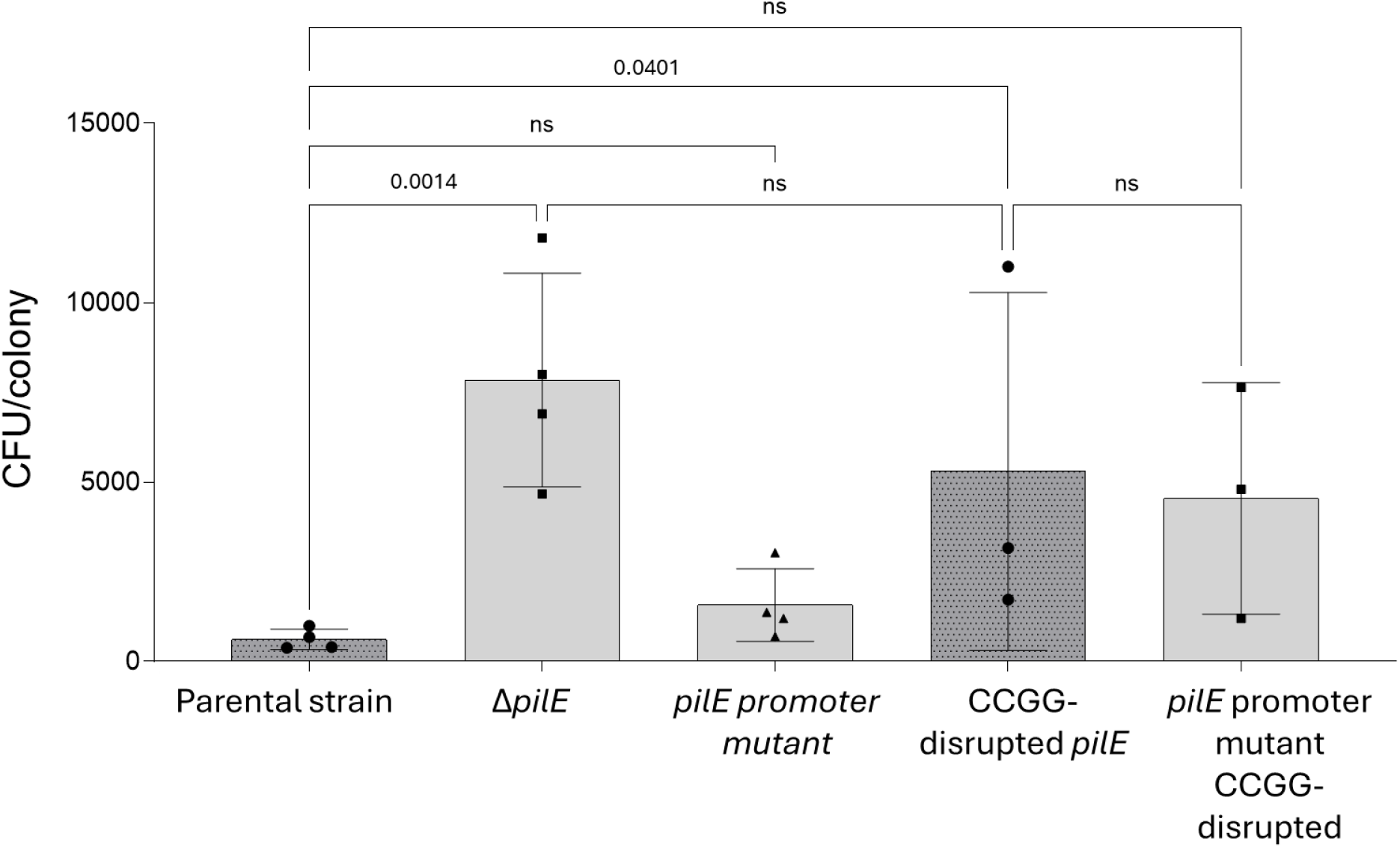
Measure of the growth of mutants carrying *pilE* mutations. CFU/colony of the strains Δ*pilE*, *pilE* promoter mutant, *pilE* CCGG-disrupted mutant and a mutant bearing both mutations at 16h of growth on GCB with IPTGMultiple comparisons were performed using Fischer’s LSD test.

**Table S3.** Strains used in this study.

| Bacterial strains |  |  |
| --- | --- | --- |
| FA1090 *recA6* | 62 | *recA6* |
| FA1090 *recA6* CCGG disrupted *pilE* | This study | SM598 |
| FA1090 *recA6* $\Delta R1.RM1$ | This study | SM323 |
| FA1090 *recA6* $\Delta R2.RM1$ | This study | SM325 |
| FA1090 *recA6* $\Delta M.RM1$ | This study | SM356 |
| FA1090 *recA6* $\Delta RM1$ | This study | SM310 |
| FA1090 *recA6* $\Delta M1.RM2$ | This study | SM362 |
| FA1090 *recA6* $\Delta M2.RM2$ | This study | SM360 |
| FA1090 *recA6* $\Delta R.RM2$ | This study | SM327 |
| FA1090 *recA6* $\Delta RM2$ | This study | SM364 |
| FA1090 *recA6* $\Delta RM1\Delta RM2$ | This study | SM500 |
| FA1090 *recA6* $\Delta R.RM3$ | This study | SM600 |
| *N-1-60 CRISPRi ngo0873 (M.RM3)* | This study | SM602 |
| FA1090 *recA6* $\Delta RM3$ | This study | SM604 |
| FA1090 *recA6* $\Delta pilS_{123678}$ hexadeleted mutant | Shaohui Yin | Q409 |
| FA1090 *recA6* $\Delta pilS_{123678}$ hexadeleted mutant KanR | This study | SM564 |
| FA1090 *recA6* hexadeleted mutant::*pilS3C1* KanR | This study | SM568 |
| FA1090 *recA6* hexadeleted mutant::CCGG-disrupted *pilS3C1*<br>KanR | This study | SM570 |
| FA1090 *recA6* hexadeleted mutant *pilE* CCGG-disrupted KanR | This study | SM582 |
| FA1090 *recA6* hexadeleted mutant::*pilS3C1 pilE* CCGG-disrupted<br>KanR | This study | SM584 |
| FA1090 *recA6* hexadeleted mutant::CCGG-disrupted<br>*pilS3C1* CCGG-disrupted *pilE* KanR | This study | SM585 |
| FA1090 *recA6* $\Delta R1.RM1$ *R1.RM1* at *iga-trpB* | This study | SM527 |
| FA1090 *recA6* $\Delta R.RM2$ *R.RM2* at *iga-trpB* | This study | SM560 |
| FA1090 *recA6* $\Delta pilE$ | <sup>63</sup> | SM648 |
| FA1090 *recA6* *pilE*-10::NheI | <sup>47</sup> | SM650 |
| FA1090 *recA6* CCGG-disrupted *pilE*-10::NheI | This study | SM655 |

